# Differential ampicillin/ceftriaxone susceptibility among diverse *Enterococcus faecalis* from infective endocarditis

**DOI:** 10.1101/2021.06.07.447474

**Authors:** Kevin J. Westbrook, Gayatri Shankar Chilambi, Hayley R. Nordstrom, Alina Iovleva, Niyati H. Shah, Chelsea E. Jones, Ellen G. Kline, Yohei Doi, Ryan K. Shields, Daria Van Tyne

## Abstract

*Enterococcus faecalis* is a leading cause of infective endocarditis (IE), especially among older patients with comorbidities. Here we investigated the genomic diversity and antimicrobial susceptibility of 33 contemporary *E. faecalis* isolates from definite or probable IE cases at the University of Pittsburgh Medical Center (UPMC) between 2018 and 2020. Isolates belonging to two multi-locus sequence types (STs), ST6 and ST179, were isolated from nearly 40% of IE patients. Both of these dominant STs carried known beta-lactam resistance-associated mutations affecting the low-affinity penicillin-binding protein 4 (PBP4). We assessed the ability of ampicillin and ceftriaxone (AC) both alone and in combination to inhibit genetically diverse *E. faecalis* IE isolates in checkerboard synergy assays and an *in vitro* one-compartment pharmacokinetic-pharmacodynamic (PK-PD) model of AC treatment. ST6 isolates as well as an isolate with a mutation in the PP2C-type protein phosphatase IreP had higher ceftriaxone MICs compared to other isolates, and showed diminished *in vitro* synergy of AC. Additionally, both ST6 and ST179 isolates exhibited regrowth after 48 hours of humanized exposures to AC. Overall, we found evidence for diminished *in vitro* AC activity among *E. faecalis* IE isolates with PBP4 and IreP mutations. This study highlights the need to evaluate alternate antibiotic combinations in clinical practice against diverse contemporary *E. faecalis* IE isolates.

## Introduction

Enterococci are commensal inhabitants of the human gastrointestinal tract (1). In the antibiotic era, *Enterococcus faecalis* has emerged as a leading cause of infection, especially in immunocompromised patients (2). The ability of *E. faecalis* to cause infection is due in part to the bacteria’s ability to tolerate a variety of stresses, including desiccation, starvation, and disinfectants (3). *E. faecalis* also possesses intrinsic resistance to a variety of antibiotics, including cephalosporins (4). While the degree of cephalosporin resistance varies between *E. faecalis* isolates, some factors that contribute to this variability include penicillin-binding proteins, two-component systems, and the RNA polymerase beta-subunit (5–8).

*E. faecalis* is a common cause of infective endocarditis (IE) (9), and the prevalence of *E. faecalis* IE is steadily increasing (10). Despite the use of *in vitro* active antibiotic combination therapy for at least six weeks, mortality from *E. faecalis* IE remains as high as 30% (11, 12). Treatment guidelines for *E. faecalis* IE include using the combination of ampicillin and gentamicin (AG) or ampicillin and ceftriaxone (AC) (13, 14). AC is a promising alternative to AG because it avoids the nephrotoxicity and the need for therapeutic drug monitoring associated with gentamicin use (15). Observational clinical studies in Europe have suggested that treatment of *E. faecalis* IE with AC is as efficacious as treatment with AG, and results in lower rates of renal toxicity (16, 17). Based upon these findings, AC is now widely used to treat *E. faecalis* IE (18). Nevertheless, *E. faecalis* IE mortality rates have not improved in decades (10). We propose that the optimal treatment approach for this disease has not yet been implemented in clinical practice.

Treatment practice for *E. faecalis* IE has largely shifted from AG to AC without investigating how genetically diverse *E. faecalis* isolates respond to AC. Prior studies documenting AC synergy have utilized a limited number of laboratory strains of *E. faecalis*, which are unlikely to represent the large genetic diversity of this species (15, 19, 20). We recently reviewed 190 *E. faecalis* IE cases from 2010-2017 at the University of Pittsburgh Medical Center (UPMC), and found higher overall 90-day mortality rates among patients treated with AC vs. AG (21% vs. 8%, *P*=0.02) (21). While baseline differences between patients influenced these findings, nonetheless it seems that for some patients, the use of AC is not optimal. In this study, we investigated the genetic and phenotypic diversity of 33 contemporary *E. faecalis* isolates collected from patients with IE. We focused on characterizing the genomic diversity of *E. faecalis* sampled from IE, identifying differences in *in vitro* AC susceptibilities, and correlating these susceptibility differences with mutations in known beta-lactam resistance-associated genes.

## Results

### Clinical presentation, diagnosis, and treatment of patients with E. *faecalis* IE

A total of 33 isolates were collected from patients diagnosed with either probable (42.4%) or definite (57.6%) IE based on modified Duke’s criteria (22, 23). The median age of the patients was 63 years (range, 24-91 years) and 79% were male (Table 1). AC combination therapy was the first-line treatment for 26 patients (78.8%). The remaining 7 patients received various combinations of cefepime, ceftaroline, daptomycin, and vancomycin. Three patients (9.1%) required alterations in their antibiotic treatment courses, and two patients (6.1%) had positive blood cultures for *E. faecalis* within three months of the initiation of antibiotic treatment. The 30-day mortality rate among patients was 27.3%, and the 90-day mortality rate was 33.3% (Table 1).

**Table 1.** Demographic and clinical features of *E. faecalis* IE patients.

| Variable | N=33 |
| --- | --- |
| Age at admission<sup>1</sup>, years | 63 (43, 72) |
| Male sex | 26 (78.8) |
| White race | 24 (72.7) |
| Definite IE<sup>2</sup> | 19 (57.6) |
| Length of Bacteremia<sup>1</sup>, hours | 66.3 (42.0, 105.5) |
| Native valve IE | 17 (51.5) |
| Prosthetic valve IE | 3 (9.1) |
| Received AC, once cultures showed *E. faecalis* bacteremia | 26 (78.8) |
| Length of hospitalization<sup>1</sup>, days | 12 (9, 31) |
| Treatment failure requiring antibiotic change | 3 (9.1) |
| Relapse, positive *E. faecalis* bacteremia within 3 months of treatment | 2 (6.1) |
| Death during hospital admission | 8 (24.2) |
| 30-day mortality | 9 (27.3) |
| 90-day mortality | 11 (33.3) |
| Readmission within 3 months, any cause<sup>3</sup> | 17 (68) |

<sup>1</sup>Median value (interquartile range) is reported. For all other variables number of patients (% of patients) is reported. AC = Ampicillin Ceftriaxone.
<sup>2</sup>Definite IE based on the modified Duke Criteria for endocarditis.
<sup>3</sup>Percent readmission was calculated for patients that survived initial hospitalization.

### Genomic diversity and biofilm formation among *E. faecalis* IE isolates

The genomes of all 33 *E. faecalis* IE isolates were sequenced and a core genome phylogeny was constructed (Table S1, Fig. 1). Isolates were found to belong to 16 distinct multi-locus sequence types (STs), with an additional three isolates belonging to STs that have not yet been defined. ST6 (n=7) and ST179 (n=6) were the most prevalent STs observed, accounting for 36% of collected isolates. Isolates belonging to ST16, ST40, and ST59 were cultured from two patients each. *E. faecalis* from all other patients belonged to unique STs.

**Figure 1.**
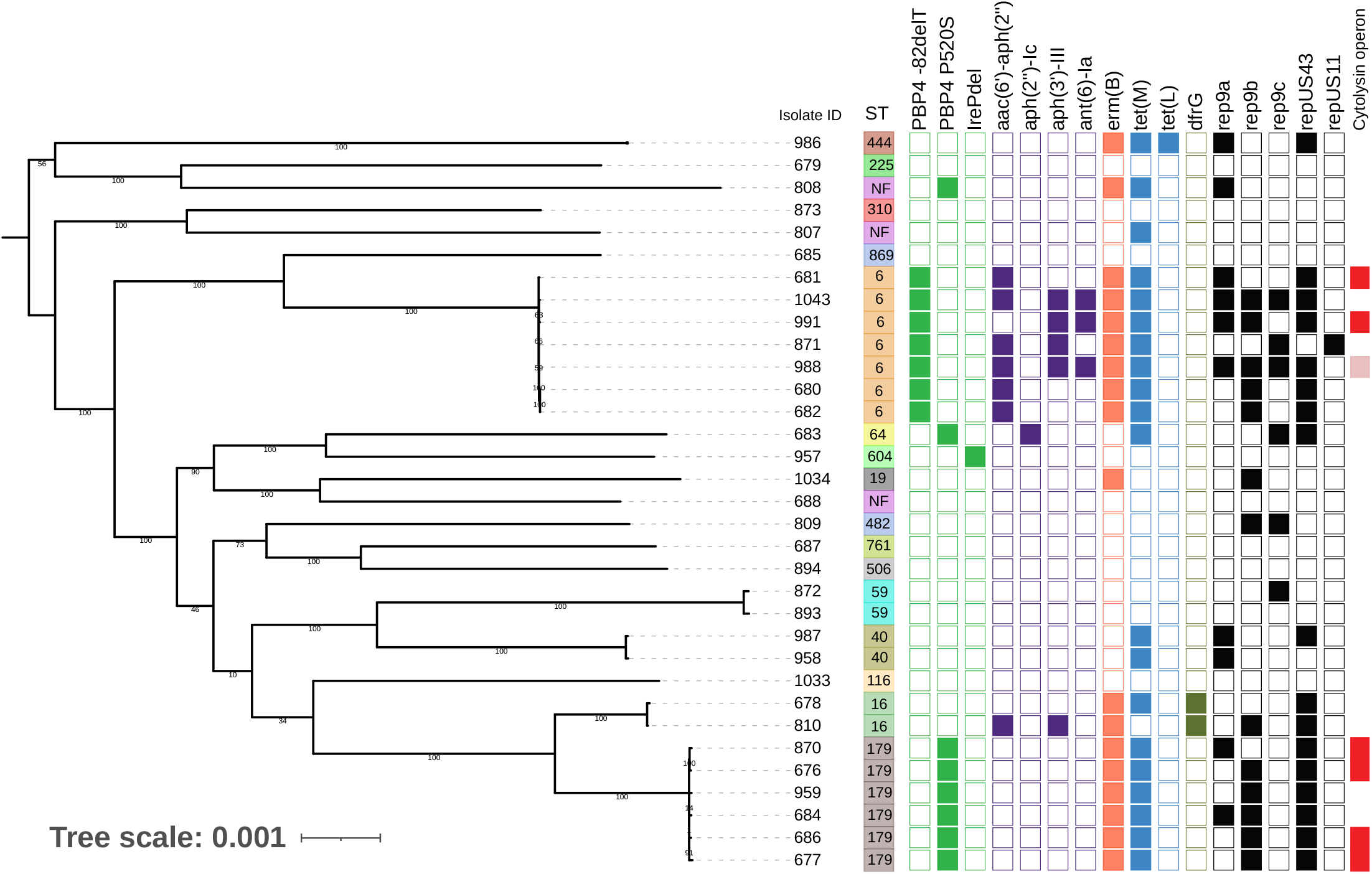
Core genome phylogeny of *E. faecalis* IE isolates. A single-copy core genome phylogeny was constructed for 33 *E. faecalis* isolates from patients with probable or definite IE. The midpoint-rooted RAxML tree was built from single nucleotide polymorphisms (SNPs) in 1980 single copy core genes identified with Roary. Tips are annotated with isolate name, multi-locus sequence type (ST), *pbp4* and *ireP* mutations, acquired drug resistance-associated genes, plasmid *rep* genes, and cytolysin operon presence (red = intact operon, pink = disrupted operon). NF = ST not found in database; *aac(6’)-aph(2”)*, *aph(2’)-Ic*, *aph(3’)-III*, and *ant(6’)-Ia* = aminoglycoside resistance (colored purple); *erm(B)* = erythromycin resistance (colored pink); *tet*(M) and *tet*(L) = tetracycline resistance (colored blue); *dfrG* = antifolate resistance (colored green).

Mutations in the low-affinity penicillin-binding protein 4 (PBP4) have been previously described in *E. faecalis*, and have been shown to affect the activity of beta-lactam agents, including cephalosporins and carbapenems (24–26). We observed PBP4 mutations in all isolates belonging to ST6 and ST179 (Fig. 1). All seven ST6 isolates carried a single base deletion 82 nucleotides upstream of *pbp4* (−82delT) while all ST179 isolates as well as two additional isolates (isolates 683 and 808) harbored a non-synonymous P520S mutation in PBP4. In addition, we identified one isolate with a 9-bp deletion in the PP2C-type protein phosphatase gene *ireP*. Mutations in *ireP* have been previously found in *E. faecalis* exhibiting high-level resistance to cephalosporins (27).

We screened all 33 *E. faecalis* genomes for acquired antimicrobial resistance genes using the ResFinder tool available through the Center for Genomic Epidemiology (CGE) (28) (Fig. 1). Aminoglycoside resistance genes were found almost exclusively in ST6 isolates (100%) and were otherwise rare in other STs (7%). All isolates belonging to ST6 and ST179 carried the erythromycin resistance gene *erm(B)* and the tetracycline resistance gene *tet*(M). The two ST16 isolates carried the antifolate resistance gene *dfrG*. All genomes were also screened for plasmid *rep* genes using PlasmidFinder (29), which identified five different *rep* genes that together were detected in 24 (70.6%) isolate genomes (Fig. 1). Among these, five genomes had only one *rep* gene identified, while the remaining 19 genomes each had 2-4 *rep* genes. ST6 and ST179 isolates carried significantly more *rep* genes than isolates belonging to non-ST6 and non-ST179 STs (ST6, mean 2.71 vs. 0.75 *rep* genes, p<0.0001; ST179, mean 2.17 vs. 0.75 *rep* genes, p=0.0007). Finally, we examined all isolates for the presence of the *E. faecalis* cytolysin, a pore-forming exotoxin that lyses bacterial and eukaryotic cells (30). An intact cytolysin operon was found in two ST6 isolates and four ST179 isolates (Fig. 1). One additional ST6 isolate had a disrupted operon. Cytolysin operons were not found in isolates belonging to any other STs.

*E. faecalis* IE is often associated with biofilm formation (31), and it is well known that biofilms protect bacteria from antibiotic-mediated killing (32). To assess differences in biofilm formation among the different *E. faecalis* IE isolates collected, we quantified the ability of each isolate to adhere to polystyrene plates using a standard *in vitro* crystal violet-based biofilm assay (33) (Fig. S1). Biofilm formation was found to be highly variable, even among isolates belonging to the same ST (Fig. S1A). Additionally, no significant differences in biofilm formation were found between isolates belonging to different STs (Fig. S1B).

### Susceptibility to ampicillin and ceftriaxone, alone and in combination, is variable among *E. faecalis* from IE

We investigated the susceptibilities of the 33 *E. faecalis* IE isolates to ampicillin and ceftriaxone by first measuring the minimum inhibitory concentration (MIC) of each isolate against each antibiotic (Fig. 2). To understand how different resistance-associated mutations in PBP4 affect antibiotic susceptibility, we separated isolates into groups based on their PBP4 genotypes. Ampicillin MICs were ≤4 μg/mL for all isolates, and did not vary significantly between PBP4 genotypes (Fig. 2A). ST6 isolates, all of which carried the PBP4 −82delT mutation, were more susceptible to ampicillin compared to other isolates, but the difference was not statistically significant (Fig. 2A). ST6 isolates were more resistant to ceftriaxone compared to both PBP4 wild type isolates (p<0.0001) as well as to isolates carrying the PBP4 P520S mutation (p=0.0006) (Fig. 2B). We also examined whether there was a correlation between ampicillin and ceftriaxone MICs by plotting them together on the same plot (Fig. 2C). We found no significant correlation between ampicillin and ceftriaxone MICs. However, the isolate with a 9-bp deletion in *ireP* had the highest ampicillin and ceftriaxone MICs among isolates lacking PBP4 mutations (Fig. 2C).

**Figure 2.**
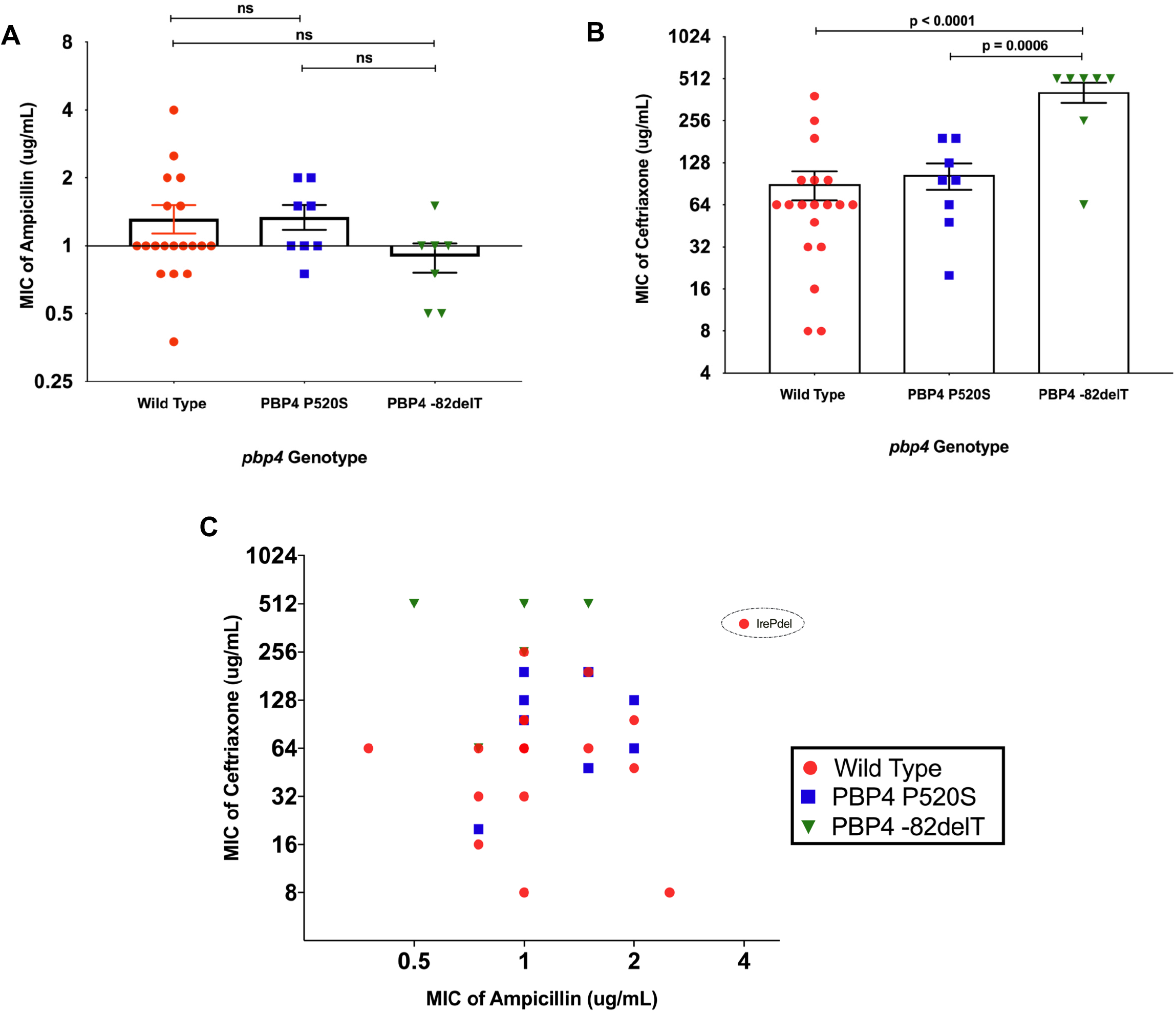
Variability in ampicillin and ceftriaxone susceptibility among *E. faecalis* from IE. Minimum inhibitory concentrations (MICs) of (A) ampicillin and (B) ceftriaxone among 33 *E. faecalis* IE isolates. MIC values were averaged between biological replicate experiments, and isolates are grouped based on their PBP4 genotype. Bars show mean MIC values and error bars show standard error of the mean. P-values are from two-tailed t-tests. ns = not significant. (C) Ceftriaxone MIC versus ampicillin MIC for each tested isolate. Isolates are colored based on their PBP4 genotypes as in (A) and (B). IrePdel refers to isolate 957, which possesses a 9-bp deletion in the PP2C-type protein phosphatase gene *ireP*.

To investigate how effectively AC could synergize to inhibit the *in vitro* growth of genetically diverse *E. faecalis* from IE, we performed standard two-antibiotic checkerboard assays on the 33 *E. faecalis* IE isolates collected (34). We found that AC demonstrated synergy against nearly all isolates, with average FIC values below 0.5 for all but three isolates (Fig. 3A). Nonetheless, FIC values among the ST6 isolates carrying the PBP4 −82delT mutation were significantly higher than both PBP4 wild type (p=0.001) and PBP4 P520S (p=0.0005) isolates. We also examined average bacterial growth across the checkerboard assay plates among isolates encoding different resistance-associated mutations (Fig. 3B). Bacterial growth was similar between PBP4 wild type and P520S isolates, while isolates encoding the −82delT mutation and the 9bp deletion in *ireP* (IrePdel) showed more growth at higher concentrations of both antibiotics. These data suggest that AC may be less active against isolates encoding *pbp4* and *ireP* mutations.

**Figure 3.**
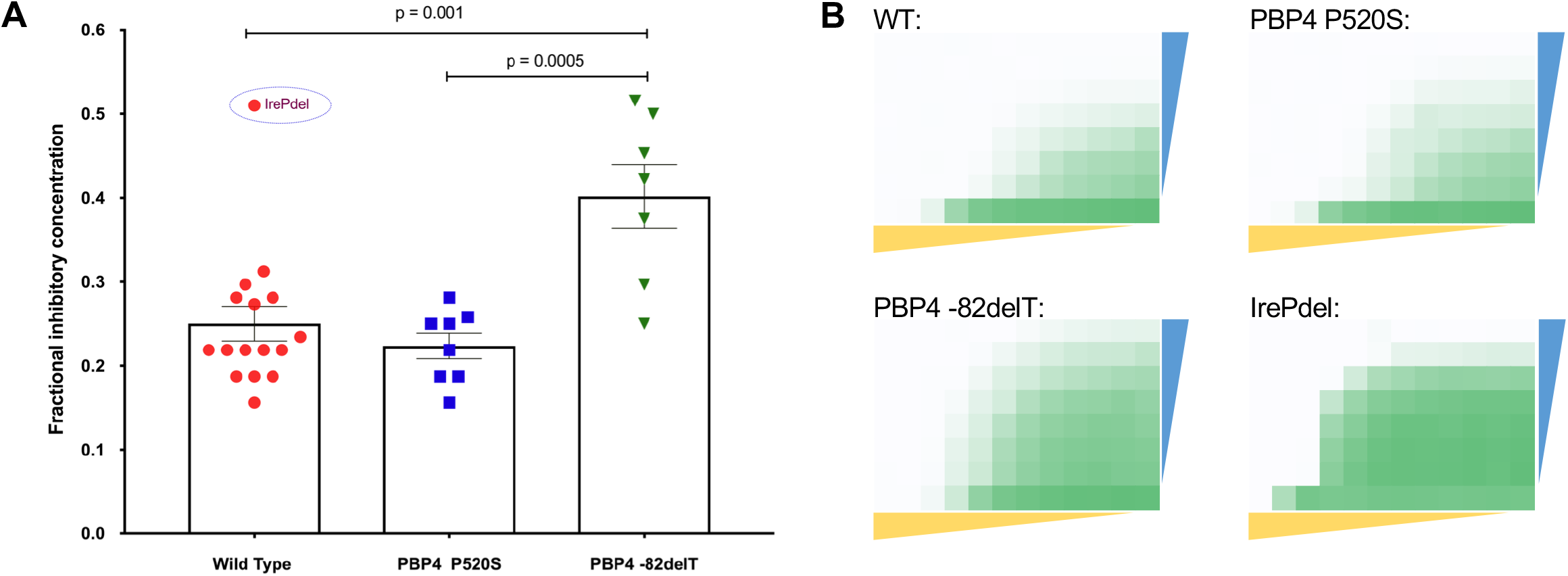
Variability in AC synergy between *E. faecalis* IE isolates. (A) FIC values calculated from AC checkerboard assays. FIC values were averaged between biological replicate experiments, and isolates are grouped based on their PBP4 genotype. Bars show mean FIC values, and error bars denote standard error of the mean. IrePdel refers to isolate 957, which possesses a 9bp deletion in *ireP*. P-values are from two-tailed t-tests. (B) Average bacterial growth in checkerboard assays. Results are separated by *pbp4* and *ireP* genotype, and growth was averaged among all assays conducted for isolates possessing each genotype. WT = wild type. Colored triangles show decreasing ampicillin (yellow) and ceftriaxone (blue) concentrations across and down each plate, respectively. The same antibiotic concentrations were tested against all isolates. Green shading indicates bacterial growth after 24 hours in each assay well, with darker shading indicating more growth.

### *In vitro* pharmacokinetic/pharmacodynamic responses to AC among *E. faecalis* from IE

Next, we tested whether diminished *in vitro* synergy observed among isolates carrying PBP4 mutations would be recapitulated in the presence of dynamic drug exposures that mimic those achieved in patients. To do so, we employed a one-compartment pharmacokinetic/pharmacodynamic (PK/PD) model to measure the killing activity and suppression of resistance following exposures to ampicillin and ceftriaxone either alone or in combination (Fig. 4). Three representative *E. faecalis* IE isolates encoding wild type PBP4 (isolate 685, ST869), PBP4 P520S (isolate 684, ST179) or PBP4 −82delT (isolate 871, ST6) were selected. Simulated antibiotic exposures were selected based on those recommended by consensus guidelines (Ampicillin 2g q 4h, Ceftriaxone 2g q 12h) (13) (Fig. 4A). Following exposure to ceftriaxone alone, isolates with PBP4 wild type or the P520S mutation showed initial killing within two hours, followed by rapid regrowth to the same levels of drug-free control arms (Fig. 4B). The isolate with the PBP4 −82delT mutation was not inhibited by ceftriaxone at any point during the experiment. By comparison, ampicillin alone resulted in rapid killing of all three isolates, with log-kills ranging from 2.48 to 4.76 log_10_ CFU/mL at 12 hours; however, each of the three isolates demonstrated regrowth after 12 hours. When administered in combination, AC was rapidly bactericidal and suppressed bacterial populations to 10^1^-10^2^ CFU/mL at 24 hours. After 24 hours, three distinct patterns were identified. Against the isolate with wild type PBP4, viable bacteria were not observed after 24 hours and bactericidal activity was maintained until the end of the experiment at 72 hours. Against the ST179 isolate encoding the PBP4 P520S mutation, late regrowth occurred, which was observed only at the final 72-hour experimental time point. Finally, against the ST6 isolate encoding the PBP4 −82delT mutation, early regrowth occurred after 24 hours (Fig. 4B). These data indicate that mutations in PBP4 decrease bacterial susceptibility to AC synergy under assay conditions and at drug concentrations that more closely reflect what the bacteria are likely to experience *in vivo* during infection.

**Figure 4.**
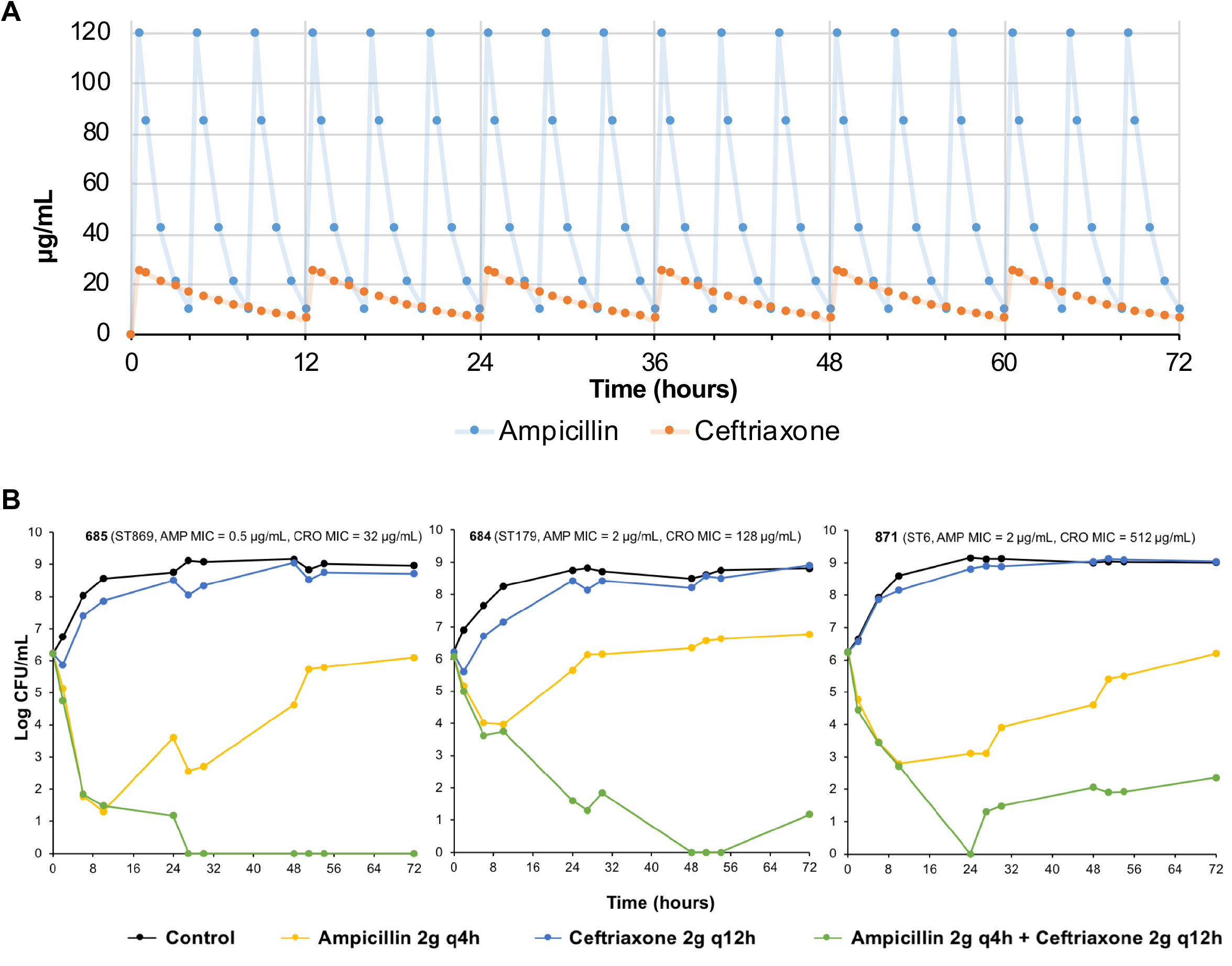
One-compartment pharmacodynamic modeling of ampicillin, ceftriaxone, and their combination against three different *E. faecalis* IE isolates belonging to different STs. (A) Simulated free serum exposures of ampicillin and ceftriaxone to achieve peak concentrations of 120 and 25.7 μg/mL, respectively. (B) Growth and killing of three different *E. faecalis* IE isolates in the presence of no drug (Control, black curves), ampicillin (AMP, yellow curves), ceftriaxone (CRO, blue curves), or ampicillin and ceftriaxone (green curves). Log_10_ CFU/mL is plotted over time for each condition. Assay limit of detection was 10 CFU/mL.

## Discussion

In this study, we characterized the genomic diversity and antibiotic susceptibility of 33 *E. faecalis* isolates from patients with probable or definite IE treated at a single medical center between 2018 and 2020. Because AC is the current front-line therapy regimen for *E. faecalis* IE, we focused on the ability of ampicillin and ceftriaxone to inhibit *E. faecalis* growth *in vitro*, using MIC and checkerboard assays as well as a one-compartment PK/PD model of bacterial killing and suppression of regrowth. Our results suggest that while AC is synergistic *in vitro* against the vast majority of *E. faecalis* IE isolates, antibiotic susceptibility levels vary widely among isolates. Notably, we observed diminished AC synergy among isolates carrying a mutation upstream of the low-affinity penicillin binding protein gene *pbp4*, as well as against an isolate with a deletion in the PP2C-type protein phosphatase gene *ireP*. We also detected bacterial regrowth within 72 hours following humanized antibiotic exposures to AC for isolates with, but not without, PBP4 mutations. These findings have potential implications for the clinical management of *E. faecalis* IE.

Among *E. faecalis* isolates collected from patients with IE, we observed abundant genetic diversity, as well as enrichment of a small number of specific genetic lineages. ST6 and ST179 were the most frequently observed lineages at our center, accounting for nearly 40% of all *E. faecalis* IE cases. This is notable because all ST6 and ST179 isolates were found to carry mutations affecting the low-affinity penicillin-binding protein gene *pbp4*, which are predicted to impact beta-lactam susceptibility (25, 26). All ST6 isolates carried a mutation 82 nucleotides upstream of *pbp4* (−82delT) that was previously shown to cause overexpression of PBP4 and increased beta-lactam resistance (25, 26). This correlated with our observation that ST6 isolates were less susceptible to ceftriaxone compared to isolates with wild type *pbp4* sequences. We also observed a non-synonymous, P520S-encoding mutation in *pbp4* among eight different isolates. In *E. faecalis*, PBP4 is a 680-amino acid membrane protein with three penicillin-binding motifs (STFK, SDN and KTG), which are located in a penicillin-binding module between amino acids 350-680 (35). The P520S substitution falls within the region between the active site-defining SDN and KTG motifs, and this mutation has been previously found to cause an increase in beta-lactam resistance (24, 25, 36). All ST179 isolates as well as two additional isolates (isolates 683 and 808) in our dataset harbored this point mutation. While we did not find significant differences in ampicillin or ceftriaxone MIC or FIC values between wild type isolates versus those with the PBP4 P520S mutation, we did observe late regrowth of the ST179 isolate that was tested in the one-compartment PK/PD experiment. This suggests that the PBP4 P520S mutation may provide increased tolerance of AC (as measured in a bacterial killing assay) without affecting overt resistance (as measured with MIC and checkerboard assays).

Most available data on AC synergy against enterococci are from *in vitro* testing (37, 38), a rabbit model of IE (15, 39), and prior clinical studies (40–43). In addition to *in vitro* susceptibility testing, we also used a one-compartment PK/PD model to monitor the efficacy of AC exposure on *E. faecalis* bacterial viability over time. This experiment is likely to more accurately reflect the *in vivo* scenario during AC treatment of patients with IE, because antibiotic exposures are modeled to simulate human dosing regimens and serum-free drug concentrations. The results of the PK/PD experiments showed that AC was rapidly bactericidal, and decreased the number of viable bacteria in all assays from 10^6^ CFU/mL to 10^2^ CFU/mL or less after 24 hours. In the case of the wild type isolate that was tested, the culture had no detectable viable bacteria after 27 hours, and viable bacteria remained below the limit of detection for the remainder of the assay. This was in contrast to the ST179 and ST6 isolates that were tested, which both showed bacterial regrowth after initial clearance of the culture. In the case of the ST179 isolate, viable bacteria were not detected after 48 hours, but they were detected again at the final, 72-hour assay time point. Regrowth of the ST6 isolate was first detected at 27 hours, and 10^2^-10^3^ CFU/mL were observed at all remaining assay time points. These data suggest that patients infected with ST6 and ST179 *E. faecalis* may experience delayed clearance of bacteremia, or recurrence of infection, compared to patients infected by *E. faecalis* with wild type PBP4 genotypes.

Prior clinical studies have shown that combination therapy with AC is non-inferior to AG therapy (16, 17). Because the use of ceftriaxone avoids the nephrotoxicity and therapeutic drug monitoring associated with gentamicin use (15), AC is now used almost universally to treat *E. faecalis* IE (18). Nearly 80% of the patients in this study were treated with AC, and no patients in this study received AG. Our findings of variable susceptibility to ceftriaxone and diminished AC synergy among a subset of isolates suggest that AC may not be the optimal treatment for all patients with *E. faecalis* IE. AG is not an ideal alternative, given similar clinical outcomes with additional toxicity. Furthermore, all ST6 isolates that we collected displayed high-level aminoglycoside resistance, meaning that gentamicin treatment would be ineffective for the same patients who may be at higher risk for delayed clearance of bacteremia or recurrent infection due to decreased AC susceptibility. Therapeutic alternatives to AC that were given to patients in this study included ceftaroline, daptomycin, and vancomycin, however these agents (and combinations of them) have not been rigorously assessed for the treatment of *E. faecalis* IE. Nonetheless, our findings suggest that alternative therapies to AC, particularly those that are effective against AC non-susceptible and AC-tolerant isolates, are needed.

This study had several limitations. Despite nearly two years of isolate collection from a tertiary medical center, we collected a relatively small number of *E. faecalis* isolates. While we hypothesize that patients with IE due to *E. faecalis* ST6 and ST179 may experience worse outcomes due to diminished AC synergy mediated by mutations in *pbp4*, our sample size is currently too small to test this hypothesis. We plan to continue to collect *E. faecalis* IE isolates from our center and other locations, and to ultimately build a cohort that is large enough to identify associations between bacterial genotypes, *in vitro* antibiotic susceptibilities, and clinical outcomes. Another potential limitation is that the checkerboard and one-compartment PK/PD assays we conducted may not accurately mirror bacterial responses to these antimicrobial combinations *in vivo*. For example, biofilm formation is likely to play a role in modulating antibiotic susceptibility during endocarditis, however in this study we did not measure how antibiotic susceptibility and biofilm formation might be linked to one another. This will be a focus of our future work, in which we also plan to employ relevant animal models of infection to assess the effectiveness of AC (as well as other alternative combinations) against genetically diverse *E. faecalis* isolates from patients with IE.

Overall, this study provides an initial description of the genomic diversity and variability in *in vitro* AC susceptibility among contemporary *E. faecalis* causing IE at a single medical center. Our analysis shows that bacteria belonging to two different STs dominate the *E. faecalis* IE population at our center. Both of these STs carry mutations that alter the *E. faecalis* PBP4, and our *in vitro* data suggest that ST6 isolates in particular may be difficult to effectively treat with AC. Further studies are needed to understand whether and how these findings impact clinical outcomes in patients with *E. faecalis* IE. Such studies could lead to improved treatment strategies for *E. faecalis* IE, which would ideally help address the high morbidity and mortality associated with these infections.

## Materials and Methods

### Collection of isolates

*E. faecalis* isolates were collected from patients with definite or probable IE at the University of Pittsburgh Medical Center between 2018 and 2020. Clinical demographics and outcomes were collected through a retrospective search of patients’ electronic medical records (EMR). Blood cultures and bacterial isolates were collected as part of routine clinical care. This study was approved by the Institutional Review Board at the University of Pittsburgh under protocol #20020039.

### Whole genome sequencing and analysis

Genomic DNA was extracted using a DNeasy Blood and Tissue Kit (Qiagen, Germantown, MD) from 1-mL bacterial cultures grown overnight in Brain Heart Infusion (BHI) media. Next-generation sequencing libraries were prepared with a Nextera XT kit (Illumina, San Diego, CA), and libraries were sequenced on an Illumina MiSeq or NextSeq using 150-bp (NextSeq) or 300-bp (MiSeq) paired-end reads. Genomes were assembled with SPAdes v3.13.0 (44), annotated with prokka (45), and were compared to one another with Roary (46). A core genome phylogenetic tree was generated using RAxML with the GTRCAT substitution model and 100 iterations (47). Sequence types, antimicrobial resistance genes, virulence factors, and plasmid replicons were identified using online tools provided by the Center for Genomic Epidemiology (https://cge.cbs.dtu.dk). Illumina read data for isolates sequenced in this study have been submitted to NCBI under BioProject PRJNA729754, with accession numbers listed in Table S1.

### Biofilm assay

Microtiter plate-based biofilm assays were performed as previously described (33). Briefly, an overnight culture of each *E. faecalis* IE isolate was diluted 100-fold into BHI broth supplemented with 0.25% glucose. 200 μL of this dilution was plated into eight replicate wells of a 96-well untreated polystyrene microtiter plate, and plates were incubated for 24 hours at 37°C under static conditions. Planktonic cells were discarded and plates were washed three times with 250 μL 1xPBS, after which adherent bacteria in each well were stained with 200 μL 0.1% crystal violet (CV) prepared in water. After incubation for 30 minutes at 4°C, wells were washed twice with 250 μL 1xPBS to remove excess stain. Plates were dried and then 250 μL of 4:1 ethanol:acetone was added to each well to solubilize the CV-stained biofilms. After incubation for 45 minutes at room temperature, the absorbance in each well was measured at 550 nm using a Synergy H1 microplate reader (Biotek, Winooski, VT). Negative control wells contained 200 μL of BHI broth supplemented with 0.25% glucose and no bacteria.

### Antimicrobial susceptibility and synergy testing

Ampicillin and ceftriaxone susceptibilities and synergy were determined by 96-well plate checkerboard assay (34). Briefly, 96-well plates were prepared containing two-fold serial dilutions of each antibiotic, either alone or in combination, in 100 μL of Mueller Hinton Broth (MHB). Overnight cultures of each *E. faecalis* isolate grown in MHB were diluted to an OD_600_ of 0.1, were then diluted 1:1000 into fresh MHB, and 100 μL of this dilution was transferred into each well of the 96-well plate, yielding approximately 10^5^ CFU per well. Plates were incubated for 24 hours at 37°C under static conditions, and growth in each well was analyzed by both visual inspection and by OD_600_ measurement using a Synergy H1 microplate reader (Biotek, Winooski, VT). Assays were conducted in biological replicates. Minimum inhibitory concentrations (MICs) were determined according to methods established by the Clinical and Laboratory Standards Institute (48). Fractional inhibitory concentrations (FICs) were calculated with the following equation: FIC = (C_A_/MIC_A_) + (C_C_/MIC_C_), where MICA and MIC_C_ are the MICs of ampicillin and ceftriaxone alone, and C_A_ and C_C_ are the concentrations of each drug in the well with the strongest inhibition. Synergy was defined as FIC<0.5. The same concentration ranges of each antibiotic were tested against each isolate. Two isolates did not yield FIC values because they did not grow in any assay wells containing both antibiotics.

### In vitro one-compartment pharmacokinetic-pharmacodynamic (PK-PD) model

A well-established one-compartment PK-PD model was employed to simulate free serum concentrations of ampicillin 2 grams (g) every 4 hours (*f*Cmax: 120 μg/mL, half-life: 1h, protein binding: 20%) and ceftriaxone 2 g every 12 hours (*f*Cmax: 25.7 μg/mL, half-life: 6h, protein binding: 90%) (13). A working volume of 250 mL of cation-adjusted Mueller-Hinton (CAMH) broth in a glass model with 4 ports (inflow, outflow, sampling, and ventilation) was used and maintained at 37°C. Peristaltic pumps controlled the flow of media to achieve desired antibiotic concentrations. In combination studies, flow rates were set to achieve ampicillin exposures given a shorter half-life and supplemental ceftriaxone was added using methods described by Blaser et al. (49). All drugs were infused over 30 minutes by programmable syringe pumps. Samples were removed at 0, 2, 6, 10, 24, 27, 30, 48, 51, 54, and 72 hours, centrifuged at 5,000 rpm for 5 minutes, washed in cold saline, and resuspended to the original volume to prevent antibiotic carryover. Each sample was then serially 10-fold diluted and plated on CAMH agar to enumerate surviving bacteria. A starting inoculum of 1×10^6^ CFU/mL was used; bactericidal activity was defined as a ≥3-log_10_ CFU/mL decrease.

### Statistics

Differences in *rep* gene abundance, antibiotic MICs, and FICs were assessed with unpaired two-tailed t-tests.

## Acknowledgements

This study was supported by the Department of Medicine at the University of Pittsburgh School of Medicine. Y.D. was supported by R01AI104895 and R21AI151362 from the National Institutes of Health. R.K.S. was supported by R03AI144636 and R21AI151363 from the National Institutes of Health. A.I. was supported by the Physician Scientist Incubator Program at the University of Pittsburgh, which is sponsored by the Burrows Wellcome Fund. The funders had no role in study design, data collection and analysis, decision to publish, or preparation of the manuscript.

---

**Figure S1.**
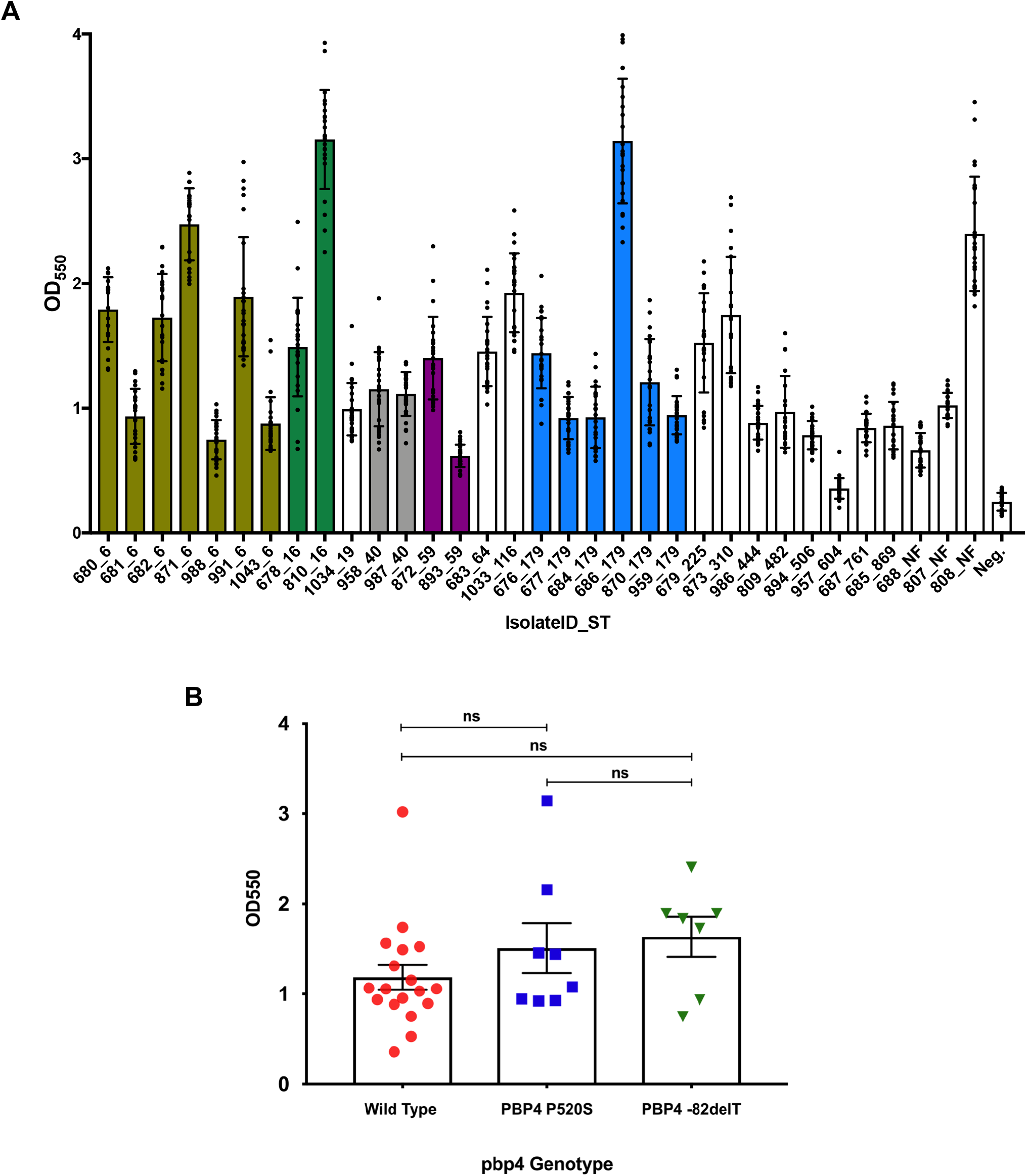
Variable biofilm production among *E. faecalis* IE isolates. (A) *In vitro* biofilm production of 33 *E. faecalis* IE isolates, organized by sequence type (ST). Biofilm formation was measured as the OD at 550nm using a standard crystal violet-based assay. Bars show mean crystal violet absorbance, and error bars show standard deviation among three biological replicate assays, each with eight technical replicates. STs with more than one isolate are colored; singleton STs are white. Neg. = negative control. (B) Biofilm formation among isolates grouped by their PBP4 genotypes. Bars show mean crystal violet absorbance values and error bars denote standard error of the mean. ns = not significant.

